# A flexible ontology for inference of emergent whole cell function from relationships between subcellular processes

**DOI:** 10.1101/112201

**Authors:** Jens Hansen, David Meretzky, Simeneh Woldesenbet, Gustavo Stolovitzky, Ravi Iyengar

## Abstract

Whole cell responses arise from coordinated interactions between diverse human gene products functioning within various pathways underlying sub-cellular processes (SCP). Lower level SCPs interact to form higher level SCPs, often in a context specific manner to give rise to whole cell function. We sought to determine if capturing such relationships enables us to describe the emergence of whole cell functions from interacting SCPs. We developed the “Molecular Biology of the Cell” ontology based on standard cell biology and biochemistry textbooks and review articles. Currently, our ontology contains 5,385 genes, 753 SCPs and 19,180 expertly curated gene-SCP associations. Our algorithm to populate the SCPs with genes enables extension of the ontology on demand and the adaption of the ontology to the continuously growing cell biological knowledge. Since whole cell responses most often arise from the coordinated activity of multiple SCPs, we developed a dynamic enrichment algorithm that flexibly predicts SCP-SCP relationships beyond the current taxonomy. This algorithm enables us to identify interactions between SCPs as a basis for higher order function in a context dependent manner, allowing us to provide a detailed description of how SCPs together can give rise to whole cell functions. We conclude that this ontology can, from omics data sets, enable the development of detailed multidimensional SCP networks for predictive modeling of emergent whole cell functions.

## Introduction

A major goal in systems biology at the cellular level is to understand how different subcellular processes (SCPs) that are organized as individual pathways or small networks^1^ are integrated to give rise to whole cell functions. This is critical for understanding how interactions between proteins (i.e. gene products) are required for whole cell physiological responses such as energy generation, action potential generation, contractility, cell movement, and secretion. Whole cell functions that form the basis for tissue and organ functions arise from coordinated activities of SCPs that are made up of multiple gene products, typically organized as pathways or small networks. SCPs interact with each other in a context specific manner; depending on the whole cell function, an SCP can interact with different sets of other SCPs. Delineating interactions between SCPs for cellular functions of interest can help us build whole cell models of mammalian cells. These models can be used to interpret and predict both cellular and possibly tissue level behaviors in a broad range of circumstances.

High-throughput experimental methods, such as transcriptomics and proteomics, enable us to identify many genes that are associated with whole cell function or change in cellular state. Computational tools like gene set enrichment analysis^2, 3^ allow for the identification of ranked lists of SCPs that are involved in the cell physiological functions being studied. For such analyses, there are a number of biological ontologies that are available. Gene Ontology (GO)^4, 5^ is arguably the most well-known ontology and is useful in associating gene lists obtained by high-throughput technologies with biological function. There are other valuable pathway based ontologies as well, such as KEGG (http://www.genome.jp/kegg/), Wikipathways (http://www.wikipathways.org) or Reactome (http://www.reactome.org/). Recently, a data-driven approach has been used to build a network-extracted ontology - NeXO^6^ - where molecular components and their hierarchical interrelationships have been defined and populated with gene products based on the network analysis of yeast protein-protein interactions.

Current ontologies provide an excellent base for the development of a new relational ontology that captures how interactions between SCPs can give rise to whole-cell functions. Such an ontology should not only contain relationships that are currently established, but also have the capability to enumerate interrelationships between SCPs that lie outside of the annotated hierarchical organization. The automated prediction of such relationships can enable systems level discovery of how higher level functions arise from interactions between lower level SCPs. Cell level physiological function often depends on the context specific relationships between SCPs that are not grouped together in a canonical taxonomy. A critical step in going from genotype to phenotype is the unambiguous delineation of interactions between SCPs that are organized to function coordinately to yield whole cell functions. In this study we sought to develop such a cell biology ontology that enables discovery of relationships between SCPs for whole cell function. Research in cell biology and biochemistry over the past fifty years has largely focused on identifying the molecular basis of cell physiology at various levels, and thus provides detailed descriptions of SCPs and their relationships. Such SCPs are typically organized in text books as pathways or small networks. These pathways or small networks can be used to describe diverse functions such as biosynthesis or degradation of amino acids or sugars, mRNA surveillance, actin cytoskeleton dynamics and signal flow within cells. We reasoned that using a well-recognized cell biology textbook could be a good starting point to develop a cellular ontology that reflects prior knowledge of how SCPs interact at multiple levels to give rise to whole cell functions. We used Molecular Biology of the Cell^7^ as our primary source to develop a cell biological ontology and called it the MBC Ontology. Additionally, we used biochemistry text books^8, 9^ and review articles (Supplementary Table S1) to develop an integrated backbone for the ontology. We populated the ontology with genes by combining text mining and statistical enrichment approaches of research articles available on PubMed. Currently, the MBC ontology contains 19,180 gene-SCP relationships arising from 753 SCPs and 5,385 genes. To allow flexible adaptation of our MBC ontology to different datasets, we developed an algorithm that allows for the identification of relationships between SCPs irrespective of whether they lie within or outside the initial taxonomy generated from the textbooks and review articles. This algorithm allows for enrichment analyses by integrating SCPs in a context specific manner. Such integration enables for the identification of those SCPs that form the basis for the whole cell function of interest.

## Results

### Generation and population of the MBC Ontology

We used the Molecular Biology of the Cell^7^, additional biochemistry textbooks^8, 9^ and review articles (Supplementary Table S1A/B) to define SCPs and arrange them in hierarchical parent-child relationships that typically span three or four levels. For example, the level-1 SCP Cytoskeleton dynamics is a child of the level-0 overall cell function process Molecular Biology of the Cell and has five level-2 children SCPs, among which is the SCP Actin filament dynamics (Fig.1A). The latter has 5 level-3 children SCPs and 6 level-4 grand children SCPs. In total our ontology consists of 29 level-1 SCPs, 125 level-2 SCPs, 487 level-3 SCPs and 111 level-4 SCPs (Fig. 1B). SCPs of the same level describe cell biological/biochemical functions of similar detail. Higher levels, i.e. levels that are closer to the level-0 overall cell function, contain SCPs that describe more general functions. Lower levels contain SCPs that describe more detailed sub-cellular functions. SCPs that have the same parent are siblings and belong to the same children set. Referring to the GO nomenclature, the vertical parent child relations in our ontology could be described using the term “part of”, a child is part of its parent.

**Fig. 1:**
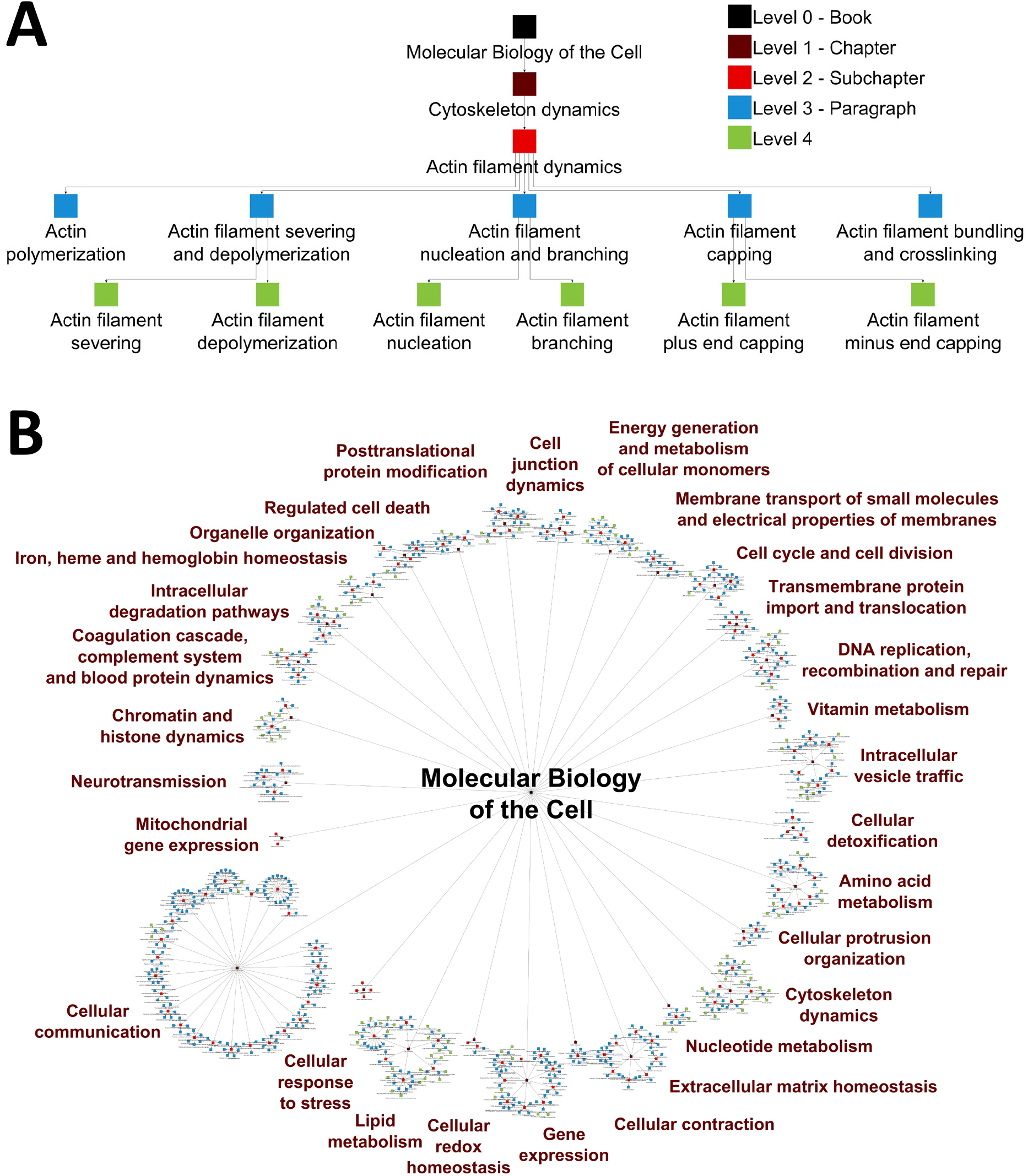
Organization of the Molecular Biology of the Cell (MBC) Ontology. (A) Example for the hierarchical organization of the MBC Ontology. The hierarchical organization of the MBC Ontology is demonstrated using the example of level-2 SCP Actin filament dynamics. It is the child of the level-1 SCP Cytoskeleton dynamics and has 5 level-3 children SCPs and 6 level-4 grandchildren SCPs. The biological detail that is described by the SCPs increases with increasing level number and corresponds to the units of the MBC textbook, where indicated. Each SCP is made of several genes/gene products that are organized as pathways or small networks. (B) Overall hierarchical organization of the MBC Ontology. The MBC Ontology currently consists of 753 SCPs that are hierarchically organized from level-1 to level 4 as shown in Fig. 1a. Processes are colored as indicated in Fig. 1A. Names describe all level-1 SCPs.

While designing the MBC hierarchy, we realized that 3 to 4 levels are largely sufficient to cover the different levels of detail that are discussed in standard cell biology text books. We did not find any advantage, especially for enrichment analyses, in defining SCPs in greater detail for functional capability than those presented in standard text books. Overall, the SCP hierarchy of the MBC ontology resembles the flatter hierarchy of the KEGG modules than the deeper hierarchy of Gene Ontology. Our aim was to design an ontology that is built upon the organization of cell biology knowledge put together by experimental scientists in standard text books and reviews. Consequently, we did not use existing ontologies to populate our ontology with genes, rather we started de novo.

We used a combination of text mining and enrichment analysis to associate genes with SCPs. We downloaded SCP-specific abstracts from the PubMed website using SCP-specific PubMed queries (Supplementary Table S1A). To identify genes and other biological entities in these abstracts, such as metabolites, we generated a dictionary for text terms of biological entities (Supplementary Fig. S1-12, Supplementary Tables S2-31). We used the dictionary to count the number of articles within each SCP-specific article set that mention a particular gene at least once (step 1 in Fig. 2 and Supplementary Fig. S13-14). The identified gene lists contained many false positive gene-SCP associations (Supplementary Fig. S15) since SCP specific PubMed abstracts do not only contain gene products that belong to that particular SCP; they may also contain gene products that interact with that SCP, or are processed by it, but are part of another SCP. Cell biologists and biochemists often select proteins to study a particular SCP that in reality belong to a different SCP, so the article counts of these proteins are normally higher in the abstract set of the SCP within which it has primary function. For example, transferrin, along with its receptor, is internalized by clathrin-mediated endocytosis and transferrin is often used to investigate the endocytosis process. In reality, the transferrin receptor belongs to the SCP Cellular iron uptake as transferrin is the major carrier of iron in blood. Similarly, many proteins that are glycosylated in the Golgi are mentioned as substrates in abstracts that describe the glycosylation machinery; however, these proteins may have different functions and hence belong to different SCPs, distinct from the SCPs describing the glycosylation pathway. To remove such false positive gene associations, we subjected the abstract counts to two rounds of statistical enrichment analysis, each followed by the automatic removal of non-selective genes, i.e. genes that belonged to a different SCP. Both times, we used Fisher’s exact test to calculate a p-value for each gene-SCP association (Supplementary Fig. S16). Afterwards, our algorithm compared the p-value distribution that was obtained for each gene and kept only those SCP associations of that gene which were associated with the most significant p-values (Supplementary Fig. S17, see methods for details). The first enrichment calculated, for each gene, the selectivity of its association with a particular SCP in comparison to all other SCPs of the same level (Fig. 2, step 2a). The second enrichment calculated the selectivity of each gene-SCP association in comparison to all other SCPs of the same children set (Fig. 2, step 3). The second enrichment step was necessary to correctly distribute the genes between the different children set processes. In many cases a gene that belonged to a particular SCP was also associated with one or more of its sibling SCPs after the first enrichment step (Supplementary Fig. S15). SCPs of the same children set often contained very high abstract counts for that gene in comparison to the other SCPs of the same level, causing the calculation of very low p-values for the association of that gene with all sibling SCPs. The direct comparison of the abstract counts between the sibling SCPs in the second enrichment step increased the accuracy of our algorithm to correctly associate the gene with that (sibling) SCP it belongs to. Gene level-2 and level-3 SCP associations were manually validated (Fig. 2, step 4, Supplementary Fig. S18) (Supplementary Text S1), and the ontology was repopulated after integration of the manual validation results (for details see methods and Fig. 2). For the population of level-1 and level-4 SCPs we used a modified protocol. Since the first enrichment step depends on a sufficiently high number of SCPs (>100) within the same level that was not the case for level-1 SCPs, as well as a broad coverage of known biology that was not the case for level-4 SCPs, we replaced the first enrichment step for these levels by an inheritance procedure (Fig. 2, step 2b). We kept only those genes in level-1 SCPs that were at least part of one of its level-2 children or level-3 grand children SCPs. Similarly, we kept only those genes of each level-4 SCP that were also part of its level-3 parent. The second enrichment step (Fig. 2, step 3) was applied to level-1 and level-4 SCPs as described above. We then added all genes of level-3 SCPs to their level-2 parent and level-1 grandparent SCPs if they were not already associated with them (Fig. 2, step 5). Similarly, we added all genes of level-2 SCPs to their level-1 parent SCPs.

**Fig. 2:**
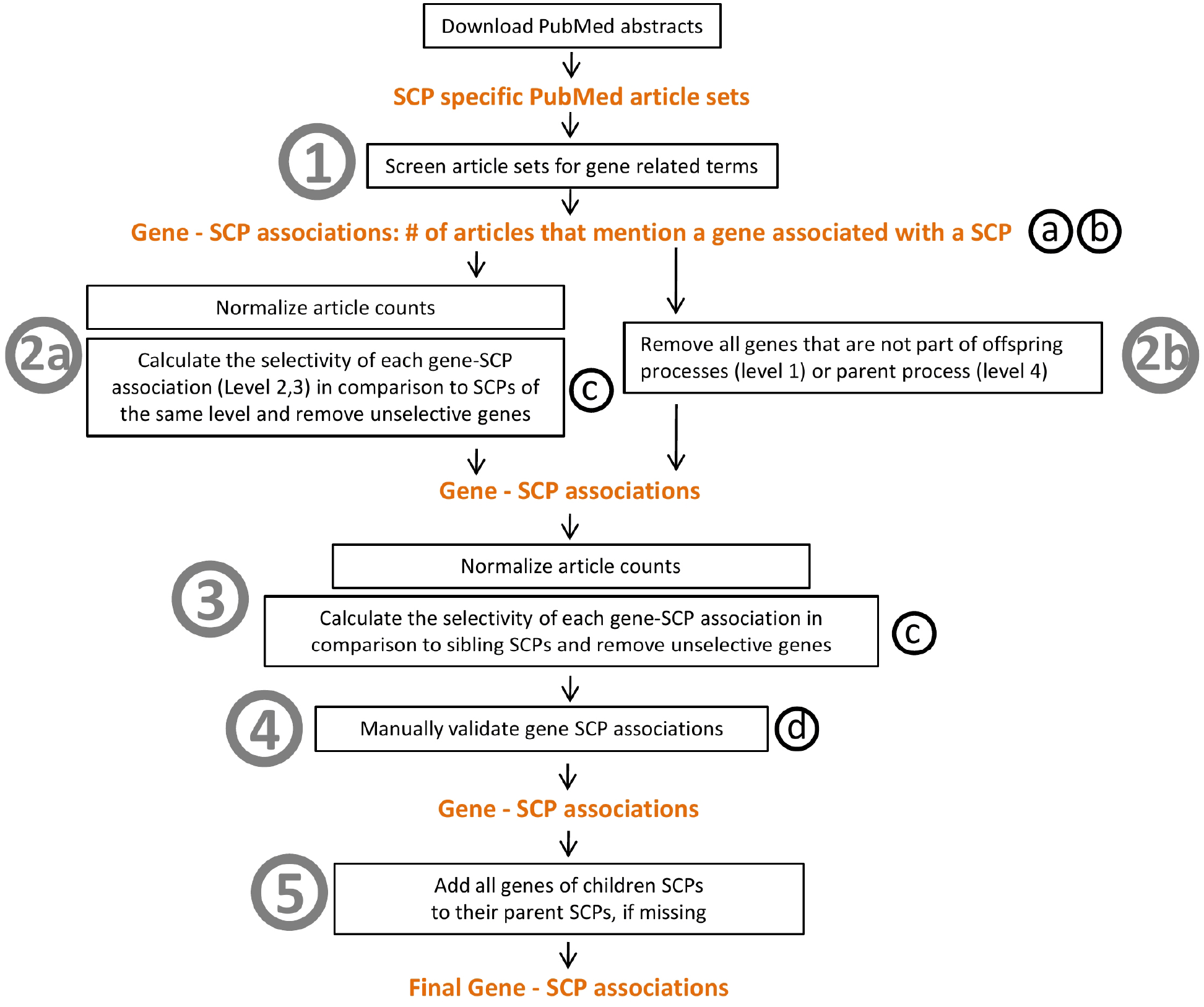
Computational pipeline to populate SCPs with genes/gene products. Circled numbers indicate steps used to populate the SCPs with genes/gene products. Circled small letters indicate the following operational modifications that were incorporated based on the manual validation of the gene-SCP associations. (a) Remove genes that are the results of misinterpreted terms during the text mining of the PubMed article sets. (b) Decrease the abstract counts of all genes that were assigned to an SCP when they belong to a sibling SCP. (c) Never remove a gene that was manually identified as a true positive in an earlier version of the MBC Ontology. (d) Remove all genes that were manually identified as false positives. See results and methods for details.

### Analysis of the MBC Ontology

Our populated ontology contained 753 SCPs and 5,385 genes organized in 19,180 gene-SCP associations. Level-1 SCPs consisted on average of 219 +/− 215 genes, level-2 SCPs of 55 +/− 47genes, level-3 SCPs of 11 +/− 11 genes and level-4 SCPs of 6 +/− 7 genes (Fig.3, Supplementary Table S32). Though the current version of the MBC Ontology is smaller than Gene Ontology or Reactome (Table 1), it is - to our knowledge - the only ontology with a clearly defined focus on cell biological function with the stated aim of covering most of the cell biology presented in standard biology text books.

**Fig. 3:**
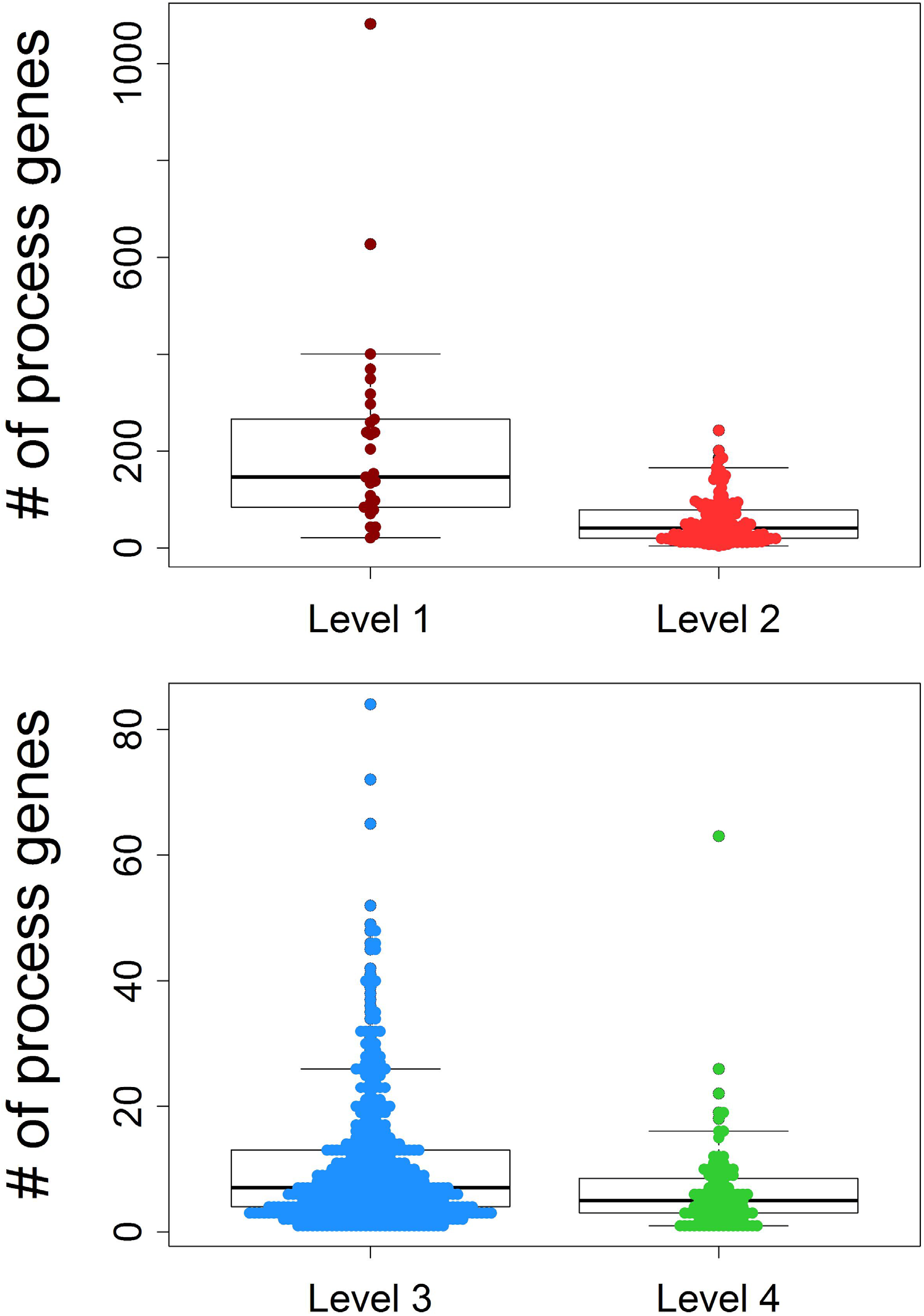
Numbers of genes per SCPs at various levels. The number of genes that were associated with each level-1 SCP and level-2 SCP as well with each level-3 and level-4 SCP was counted and visualized. Each colored dot corresponds to the number of genes of one SCP of the indicated level.

**Table 1:** Comparison of the MBC ontology with other ontologies. For each of the indicated ontologies we counted the number of processes, genes and gene process associations. In its current version, the MBC ontology ranks in the middle of the selected ontologies. A main difference of the MBC ontology with the other ontologies is its strict focus on cell biology and its aim to cover most of the cell biology that is presented in cell biological text books.

| **Ontology** | **Processes** | **Genes** | **Process-gene associations** |
| --- | --- | --- | --- |
| Gene Ontology biologicalprocess 2017 | 5192 | 14264 | 301878 |
| Reactome 2016 | 1530 | 8973 | 97479 |
| Molecular Biology of the Cell | 753 | 5385 | 19180 |
| Kegg 2016 | 293 | 7010 | 25391 |
| Wikipathways 2016 | 281 | 4950 | 12853 |
| Biocarta 2016 | 237 | 1348 | 4420 |

We were interested in whether our population algorithm was able to generate consistent results in vertical relationships between level-2 parent and level-3 children SCPs, i.e. if our ontology is internally consistent. If the hierarchical (i.e. vertical) SCP relationships reflect common biological understanding, there should be a significant overlap between the genes of parent and children SCPs. For this analysis, we focused on level-2 and level-3 SCPs that were populated based on both enrichment steps, and considered their gene composition before the addition of the genes of children SCPs to their parent SCPs. We determined for each level-3 child SCP how many of its genes are also associated with its level-2 parent SCP (Supplementary Fig. S19A, Supplementary Table S33). The average percentage of child SCP genes that were also associated with its annotated parent was 45.9% +/− 36.9% (median: 38.5%). To document the accuracy of our annotation, we identified for each level-3 child SCP that level-2 SCP that contained the highest number of the genes of the level-3 child SCP (Supplementary Fig S19B). For 498 level-3 SCPs the identified best matching level-2 SCP was the annotated parent SCP, for 28 level-3 SCPs we identified more than one level-2 SCPs that contained the annotated parent SCP, while for 184 level-3 SCPs the identified level-2 SCP was not the annotated parent SCP.

To ascertain a (nearly) complete coverage of the sub-functions of a level-2 parent SCP by its level-3 children SCPs, we determined how many genes of a parent SCP were also part of at least one of its children SCPs (Supplementary Fig. S19C). On average the combined children SCPs contained 76% of the parental genes. To analyze if the children SCPs describe mutually exclusive functions, we documented the overlap between the parent genes and the union of all its children genes after the removal of one or more children SCPs. If every child SCP describes a unique sub-function of the parental SCP, the removal of every child SCP should decrease the number of overlapping children set genes that are part of the parent SCP. For one of the level-2 parent SCPs at a time, we sequentially removed one to all of its level-3 children SCPs, re-populated the remaining level-3 SCPs using the entire pipeline for population of SCPs (Fig. 2, excluding step 4 (manual validation)). We re-calculated the overlap between the genes of the parent and remaining children SCPs (Supplementary Fig. S18C), and observed a continuous decrease in the number of overlapping genes. This further supports the internal consistency of our ontology and the biological correctness of our annotated hierarchy.

### Genes within an SCP show correlated expression

Genes that are involved in the same sub-cellular function act in concert with each other and could correlate in their gene expression over different conditions or tissues^10^. We analyzed if our algorithm populates SCPs with genes that show high co-expression and can therefore be assumed to be functionally related. Using genome-wide expression data, measured in different human tissues (http://www.gtexportal.org)^11^, we calculated the Pearson correlation between all pairs of genes. For each level, we then distributed the correlations into two groups: the first group contained all correlations between any two genes that were associated with at least one common SCP of that level, the second group contained all correlations between any two genes that did not belong to any common SCP. In this analysis, we focused on level-3 SCPs since they describe clearly defined sub-cellular functions and do not summarize multiple different or opposite sub-cellular functions as is the case for many level-1 and level-2 SCPs. The correlations between the genes of both groups belong to two significantly different populations (Fig.4), providing evidence that genes of the same level-3 SCP show more correlated expression. As expected, the two populations for level-1 and level-2 SCPs were less different (Supplementary Fig. S20). We conclude that the gene co-expression analysis further supports the ability of our algorithm to populate SCPs with genes that are most likely part of the function of the SCP.

**Fig. 4:**
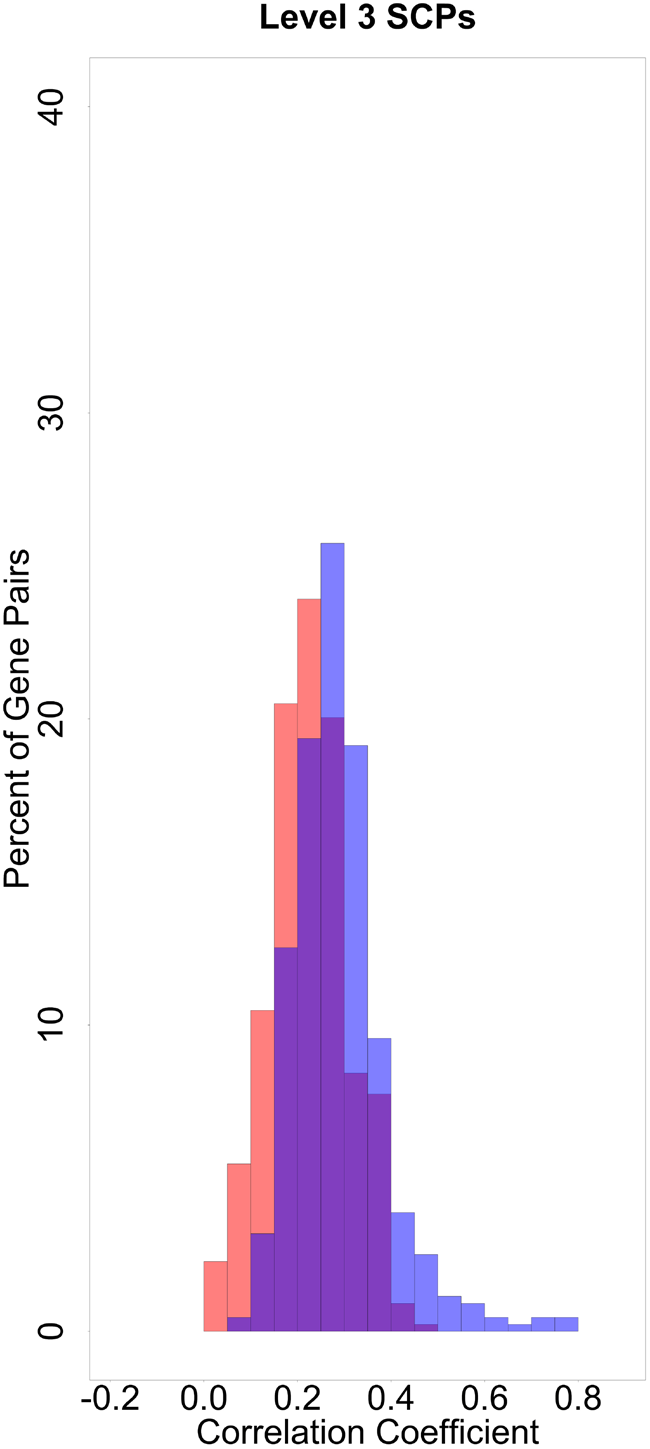
Higher levels of co-expression in various tissues for genes that are part of the same level-3 SCPs. We determined the Pearson correlation coefficient between any two genes that belong to level-3 SCPs for their expression in 30 different tissues using the GTEx data. We distributed the correlations into two groups: The first group (blue) contained all correlated gene pairs that were part of at least one level-3 SCP (15,024 pairs), the second group (red) all other correlated gene pairs (6,719,761 pairs). For ease of visualization both groups were normalized to 100 and percentage of pairs for the various correlation coefficients are shown. The Kolmogorov-Smirnov test shows that gene expression correlations significantly differ between the two groups (p-value = 4.885e-15).

### Inference of relationships between SCPs at the same level

The competitive nature of our algorithm to populate SCPs allowed us to infer new SCP-SCP (horizontal) relationships between level-3 SCPs independent of their annotated family hierarchical (vertical) relationships. These multidimensional inferred relationships can be a guide for biological understanding, as well as form the basis for a dynamic enrichment analysis that considers context specific SCP interactions. SCPs of the same level compete with each other for genes during the population approach. The degree of competition between any pair of two SCPs can be used to define weighted horizontal SCP-SCP relationships: the more two SCPs compete with each other for genes, the more they are related to each other. To identify the degree of competition between any two SCPs of the same level, we performed leave-one-out analysis. We iteratively removed one SCP at a time from the level-3 SCP set, and repopulated the remaining level-3 SCPs as described above (Fig. 2). For each of the remaining SCPs, we compared the alternative gene sets with the original gene sets to identify those remaining processes that were influenced by the removal of the current SCP. We developed an additional algorithm to quantify these effects. This algorithm considers new gene additions to the remaining SCPs as well as rank increases of original genes within the remaining SCPs. This way we could generate a level-3 SCP-SCP interaction network where edges link the removed SCPs to the remaining SCPs and the edge width is proportional to the distribution of genes between these two SCPs. We predicted 2507 interactions between level-3 SCPs that connect SCPs within and beyond the annotated families (see Fig. 5, Suppl. Table S35). Though we can readily assign a description to the annotated interactions between child and parent SCPs, the nature of the predicted relationships are case dependent. Examples for such relationships are “regulates” (e.g. HIF1 signaling regulates glycolysis and gluconeogenesis), “depends on” (e.g. Electron transport chain depends on Riboflavin metabolism) and “interacts” (e.g. Urea cycle interacts with Aspartate and arginine metabolism).

**Fig. 5:**
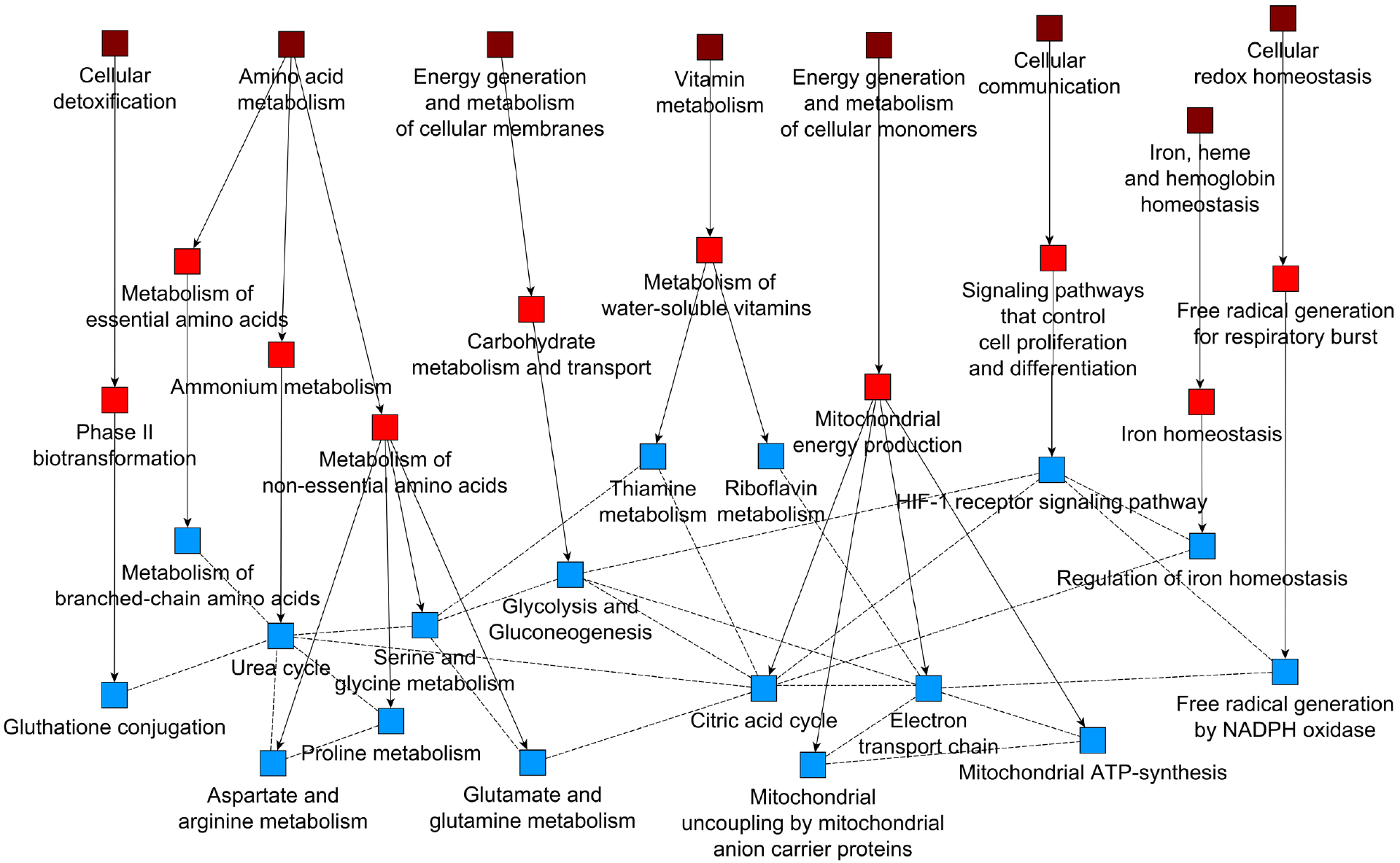
Algorithm to connect level-3 SCPs beyond family relationships defined by the initial taxonomy. One level-3 SCP at a time was removed from the ontology and the remaining level-3 SCPs were re-populated. Each of the remaining SCPs was analyzed, if it contained genes of the removed SCP, either as a new member or as a member that showed increase in rank. Changes were quantified and results associated with SCP pairs. The newly identified SCP interactions are shown by dotted lines while the original taxonomy is shown as solid arrows. All predicted interaction partners of three SCPs Urea cycle, Citric acid cycle, and Electron transport chain are shown. To illustrate the hierarchical relationships all level-2 parent SCPs (light red) and level-1 grandparent SCPs (dark red) are also shown.

### Dynamic enrichment analysis enables the emergence of whole cell functions from interactions between SCPs

To test if our ontology can be used to describe whole cell functions as sets of interacting SCPs, we used data from three published studies, using different high-throughput approaches to characterize a well-defined whole cell phenotype. We wanted to determine if we could predict the phenotype described in the publication from the results of our enrichment analysis without any prior assumptions. We first analyzed the data using standard enrichment analysis, and then compared this approach to a new type of enrichment analysis that considers context specific interactions between SCPs, which we call dynamic enrichment analysis. Standard enrichment analysis uses statistical tests such as Fisher’s exact test to identify SCPs that are significantly associated with the experimental gene list. SCPs are then ranked by significance, followed by manual inspection and selection of those processes that can be related to the whole cell function of interest. Often the researcher focuses on the top ranked SCPs and ignores the rest. Whole cell functions are based on various SCPs which are distributed over different families and therefore are described in different sub-chapters of text books, but can function in an integrated manner. In such a scenario, SCPs with low statistical significance could still make an important contribution to whole cell function, if they interact with each other as part of a higher level context specific SCP. We used the inferred horizontal relationships between level-3 SCPs to address this question by use of the dynamic enrichment approach. For each dataset, we generated new context-specific higher level SCPs by merging two or three level-3 SCPs that were connected by inferred relationships (i.e. that were among the top 25% of the predicted level-3 SCP relationships) and contained at least one experimentally determined gene (e.g. a differentially expressed gene or a differential protein binding partner). We added these context specific level-3 SCP-units to the standard SCPs and repeated the enrichment analysis. The SCP networks obtained by the dynamic enrichment analysis described the emergence of the whole cell functions that were analyzed in the published papers used as case studies. This could be done without any prior assumptions or manual selection. Three examples are described below.

In the first case study, we tested our ontology on proteomic data by analyzing the mutant cystic fibrosis transmembrane conductance regulator (dF508 CFTR) interactome^12^. The mutated anion channel is the cause of cystic fibrosis. Impaired dF508 CFTR folding leads to its premature intracellular degradation and consequently reduces CFTR activity at the plasma membrane of bronchial epithelial cells, characterizing cystic fibrosis as a protein folding disease. Here our focus is not the disease, rather we sought to determine if we could identify the SCPs that enable protein proofreading and degradation in cells from the proteomic data without any prior assumptions. The authors investigated the different protein interaction partners of WT CFTR vs dF508 CFTR via co-immunoprecipitation and proteomic analysis. Standard enrichment analysis of the identified different interaction partners revealed SCPs that were distributed over a wide range of cellular functions: actin cytoskeleton, vesicular traffic, glucose and lipid metabolism, ribonucleoprotein biogenesis, protein folding, and quality control in the ER as well as proteasomal degradation (Supplementary Fig. S21A and B). All these processes were described by the authors after manual inspection of the genes and were thought to contribute to the reduction in the level of the mutant dF508 CFTR in the plasma membrane. Based on the statistical significance of the SCPs the results of the standard enrichment analysis would propose potential changes in ribosome metabolism, sugar metabolism and actin cytoskeleton as major mechanisms for this deficiency. Similarly, enrichment results obtained with Gene Ontology did also not allow a clear identification of the investigated phenotype (Supplementary Fig. S21C). In contrast, the dynamic enrichment analysis using the MBC ontology predicted protein folding defects and proteasomal degradation, as the major mechanism (Fig.6A, Supplementary Fig.S21D/E) for the decreased levels of the mutant CFTR protein in the plasma membrane. We summarized these SCPs under a new inferred context-specific parent SCP that we called “Protein proofreading and degradation”, allowing us to provide a logic for the hierarchical organization of SCPs that give rise to the observed cellular phenotype. It should be noted that the SCP Protein proof reading and degradation which is not in the starting taxonomy can be related to the two level-1 SCPs Intracellular degradation and Post-translational protein modification (dashed lines in Fig. 6A).

**Fig. 6:**
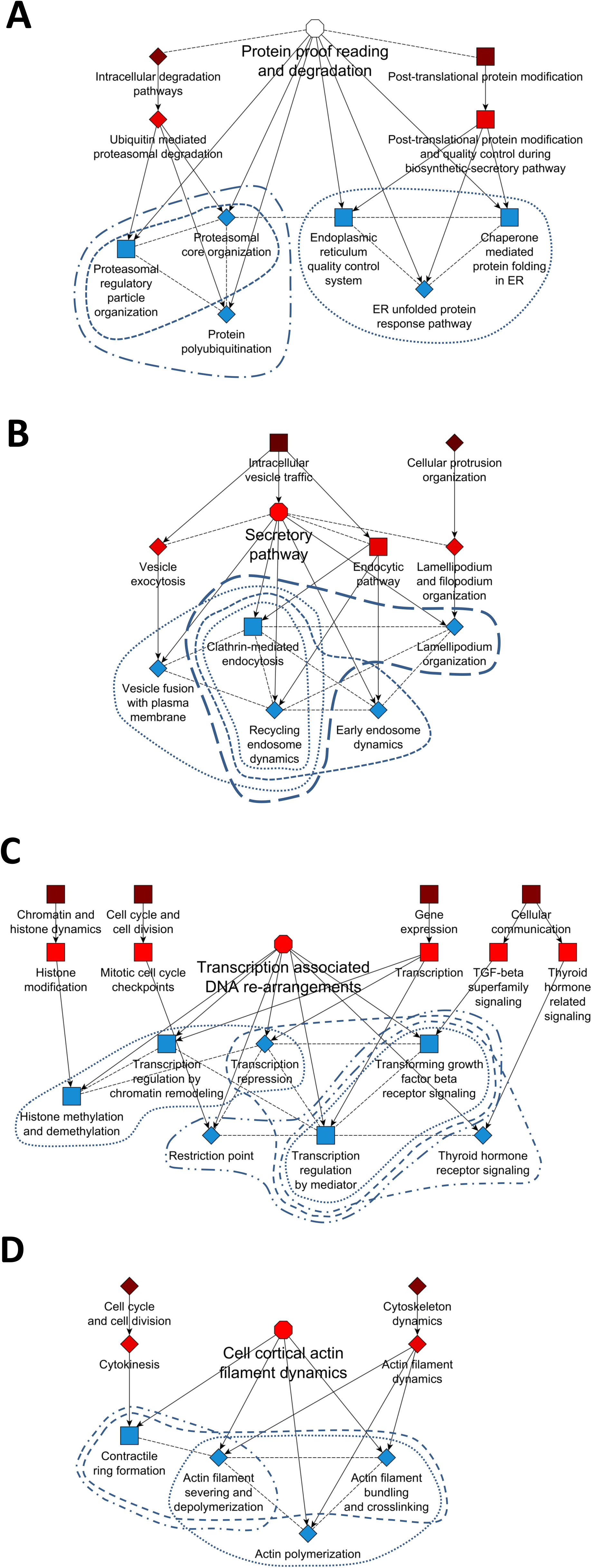
Dynamic enrichment analysis identifies context relevant higher level SCPs emerging from interactions of level-3 SCPs. Three case studies using experimental high throughput datasets from literature are used to demonstrate the ability of the MBC Ontology to identify SCPs relationships that give rise to the whole cell function that was studied in each case. SCP units were generated by merging any two or three level-3 SCPs that were related to each other based on the top 25% predicted SCP relationships and contained at least one experimentally perturbed gene. The merged SCPs were added to the original level-3 SCPs and the experimental data was analyzed for enrichment analysis using the extended set of level-3 SCPs and Fisher’s exact test. All level-3 SCPs (blue rectangles and diamonds) that were among the top 5 predictions (either as a single SCP or as an SCP unit) were connected with each other based on the predicted SCP-SCP relationships (dashed black lines). The largest SCP network was kept and assigned to a context specific higher level SCP (open [unspecified level] or light red [level-2] octagon). Annotated level-1 grandparents (dark red rectangles) and level-2 parents (light red rectangles) were added to demonstrate that the combined level-3 SCPs belong to different SCP families. Arrows connect parent SCPs with their children SCPs. Blue lines encircle level-3 SCPs that were identified as part of a new merged SCP unit. Squares indicate level-1 to level-3 SCPs that were identified by the original taxonomy. Diamonds indicate level-1 to level-3 SCPs that were only identified by the dynamic enrichment approach that enables extension of the original taxonomy. (A) Dynamic enrichment analysis identifies the inferred context relevant higher-level SCP Protein proofreading and degradation as the major disease mechanism that reduces CFTR activity at the plasma membrane in cystic fibrosis. (B) The two major parts of the Secretory pathway were identified as a major requirement for secretion. (C) Transcription associated DNA re-arrangements were identified as a major mechanism by which erlotinib increases sensitivity of breast cancer cells to doxycycline. In this case including the top 26% (instead of 25%) predicted SCP relationships resulted in the identification of a larger SCP-network that is related to the identified whole-cell function (at a 25% threshold the network contained the SCP “Focal adhesion organization” that was connected to the SCP “Thyroid hormone receptor signaling” instead of the SCPs “Histone methylation and demythelation” and “Transcription regulation by chromatin remodeling”). (D) Cell cortical actin filament dynamics were predicted to be responsible for reduction in colony formation activity of erlotinib treated breast cancer cells. See also Supplementary Fig. S21 D/E, 22 D/E and 23 D/H/I.

In the second case study, we tested our ontology using a list of genes that were identified in a genomewide RNAi screening as stimulators of the secretory pathway^13^. We sought to determine if we could identify secretory pathways as the whole cell functions from the lists of knocked out genes. The authors identified these genes by documenting the effect of their knockdown on the surface arrival of the fluorescently labeled viral protein tsO45G. Standard enrichment analysis identified hierarchically connected level-1 to level-4 SCPs that describe vesicular transport, nucleotide metabolism and gene expression (Supplementary Fig S22A and B). Standard enrichment analysis using Gene Ontology also predicted various processes in different areas of cell biology (Supplementary Fig. S22C). To identify the primary SCPs involved we conducted dynamic enrichment analysis using the MBC Ontology. The results of the dynamic enrichment analysis showed a cluster of SCPs that described actin assisted vesicle fusion and membrane recycling processes at the plasma membrane (Fig. 6B, Supplementary Fig. S22D and E), indicating that the investigated phenotype is the two major parts of the secretory pathway. Thus dynamic enrichment analysis with the MBC Ontology allows us to correctly identify the SCPs driving the studied whole cell function.

In the third case study, we used gene expression data as the test dataset. The pretreatment of the breast cancer cell line BT-20 with the EGFR inhibitor erlotinib for 4 to 24h dramatically sensitizes the cell line towards doxirubicin induced apoptosis^14^. To identify possible mechanisms for this observation, the authors analyzed differentially expressed genes (DEGs) after 6h and 24h erlotinib treatment and identified significant changes in 16 out of 34 GeneGo cellular networks that include pathways related to apoptosis and DNA damage response. They also demonstrated that treatment with erlotinib diminishes the ability of the cells to form colonies in soft agar, a standard test for metastatic potential. We sought to determine if the MBC Ontology, using the transcriptomic data could identify the SCPs by which erlotinib pretreatment sensitizes the cells to doxyrubicin. Using our ontology, we re-analyzed the DEGs. Using standard enrichment analysis we identified multiple different SCPs based on the DEGs after 6h erlotinib treatment (e.g. Actin filament dynamics, TGF-beta superfamily signaling, RNA surveillance and degradation or Pre-mRNA 3-end-cleavage and polyadenylation) (Supplementary Fig. S23A and B), as well as after 24h treatment (e.g. Apoptosis, JAK/STAT signaling pathway or Hemidesmosome organization) (Supplementary Fig.S23E/F). Among the multiple predictions that we obtained based on Gene Ontology (Supplementary Fig S23C and G) were the processes Transcription from RNA polymerase II promoter (rank 2, 6h erlotinib) and Chromatin modification (rank 8, 6h erlotinib) that propose mechanisms that could be triggered by erlotinib to sensitize the cells to doxyrubicin toxicity. We investigated, if our dynamic enrichment analysis generates results that identify a selective mechanism. Dynamic enrichment analysis of the DEGs after 6h identified a cluster of SCPs that describes transcription associated DNA rearrangements (Fig. 6C, Supplementary Fig. S23D and E), an under-recognized cellular process^15^. This SCP refers to the mechanism of doxyrubicin action involving Topoisomerase II poisoning, DNA adduct formation, and DNA intercalation that can influence DNA topology and nucleosome dynamics^16^. In agreement with two of the top 10 predictions that we obtained using Gene Ontology, the results of the dynamic enrichment analysis therefore indicate that erlotinib pre-treatment re-wires the cellular machinery at doxyrubicin’s site of action, suggesting a mechanism for increased sensitivity towards doxyrubicin that was not discussed by the authors. The dynamic enrichment results of the DEGs after 24h erlotinib treatment offer an explanation for the diminished colony formation activity induced by erlotinib. We identified a cluster of SCPs describing cell cortical actin filament dynamics (Fig. 6D, Supplementary Fig. S23H and I). Actin filament remodeling has been documented as a major cause for significant decrease in migration and colony formation^17^.

## Discussion

To understand how genomic and proteomic determinants are integrated into cell, tissue, and organ level responses, we need to understand how these determinants are organized to execute specific whole cell functions. Many individual pathways and small networks within cells combine to produce cellular phenotypes. Without an ontology that focuses on cell biological pathways and networks it will be impossible to relate genomic and proteomic characteristics to cell physiology as the basis for tissue and organ level function. The Molecular Biology of the Cell Ontology is designed to focus on sub-cellular activities that together give rise to whole cell function. For this we used the organizational logic that has been organically developed by experimental biochemists, cell biologist and physiologists. This logic is evident in textbooks and review articles and we used the organization of a standard cell biology textbook as a the basis for our organization. This specification is a major difference between the MBC ontology and other ontologies that contain biological processes at multiple levels of organization, not only relating to (a subset of) cell biology, but also to tissue/organ and organismal functions as well as diseases. Currently, the ontology contains 753 SCPs that cover a wide range of cell biological mechanisms, although the current list of SCPs is not comprehensive. Testing our ontology on varied published data sets obtained from diverse high-throughput technologies, we find that the MBC Ontology provides a reasonable organizational map that allows for intuitive understanding of hierarchical relationships for the emergence of whole cell functions.

Different whole cell functions may engage sets of similar and different pathways (i.e. SCPs), in different combinations. We developed an algorithm that enables the prediction of horizontal relationships between level-3 SCPs that are within as well as beyond the relationships between annotated sibling SCPs that are defined by vertical parent-child relationships. We used these predictions to introduce - to our knowledge for the first time - the concept of dynamic enrichment analysis that considers process dependencies in a context specific manner. The lack of consideration of such process dependencies is one of the major limitations of standard enrichment analysis^18^. By applying dynamic enrichment analysis to the test datasets, we were able to accurately predict the whole cell response being studied. The dynamic enrichment approach seems to increase the predictive power based on existing SCPs that could only be achieved by manual definition, population and validation of new SCPs otherwise. The results of our proof-of-principles studies indicate that dynamic enrichment analysis is a valuable addition to standard enrichment analysis, since the consideration of SCP dependencies enables the identification of appropriate higher level subcellular function that shows an enrichment of experimentally observed genes (e.g. DEGs). Dynamic enrichment analysis depends on the description of functional interactions between SCPs. Rigidly annotated SCP relationships are not enough for such an approach, since whole cell responses or cellular changes of state are not restricted to the borders of human made taxonomy and classification. To our knowledge such functional interactions between SCPs within and beyond annotated family relationships so far has been identified for 131 GO terms encompassing 71 interactions that are based on additional experimental data^19^. Additional interactions between GO terms based on new experimental data that cover a wide range of biological functions could also form the basis for dynamic enrichment analysis based on Gene Ontology and other ontologies.

Currently, our ontology accounts for about a quarter of the human genome and about 40% of the 13,762 genes that our text mining algorithm identified in at least one of the screened abstracts. We currently use a minimum of four/three/two/one publications for a gene to be associated with a level-1/2/3/4 SCP.

Lowering these thresholds might result in the addition of further true positive gene-SCP associations and increase the total number of genes in our ontology. Additionally, our population and validation approach allows for extension of the ontology as needed. The MBC Ontology can be easily extended beyond what we described here for additional cellular, tissue, and organ level functions. As biological validation of a function of a gene within a sub-cellular process is likely to come from detailed studies, capturing such validation in an ontology will require extensive literature study. This is essentially a big data problem wherein one requires a relatively facile approach. As part of developing the MBC Ontology, we have devised an automated text-mining algorithm that allows for a relatively rapid manual validation of functional associations. Thus we are able to combine the speed of computer searches with expert human knowledge. With appropriate training and supervision by a domain expert a single person can curate 1000 gene-SCP associations within 3-6 hours. As cell biology, biochemistry, and physiology are vast fields with different domains of expertise, this combination of computer searches and human validation can be used in the longer term to produce a definitive ontology of all sub-cellular processes that accounts for all of the gene products encoded by the human genome.

## Methods

### Generation of the hierarchical structure of MBC Ontology

The SCPs and their hierarchical organization were designed based on three standard cell biological and biochemistry textbooks^7–9^. Additionally we needed a number of review articles (Supplementary Table S1A). Based on the degree of detail, SCPs were assigned to different levels: SCPs from lower number levels were more general, while SCPs from higher number levels were more detailed. Level-0 consists of only one overall function (“Molecular Biology of the Cell”) that summarizes all whole cell functions and resembles the book. Level-0 “Molecular Biology of the Cell” is the parent of all level-1 children SCPs that resemble book chapters (e.g. “Cytoskeleton dynamics”). Each level-1 SCP has level-2 children SCPs that refer to the corresponding subchapters. Level-2 SCPs are the parents of level-3 SCPs and some level-3 SCPs have level-4 children SCPs. We use the terms same children set or siblings to refer to all those processes that have the same parent, while the term same grandchildren set summarizes all processes that have the same grandparent. The term family is used to indicate a nuclear family, i.e. it consists of a parent and its children, excluding the grandchildren.

### Populating the SCPs with genes/gene products

Our aim was to populate the SCPs with genes (i.e. NCBI gene symbols) that we identified in SCP-specific PubMed article sets. For each SCP we downloaded all titles and abstracts from the PubMed website that were obtained by an SCP specific query (Supplementary Table S1A). Authors of cell biological, physiological and biochemical articles often do not use the NCBI gene symbols, but gene synonyms, gene names, alternative gene names, names of protein complexes, and protein families. To be able to identify these terms in our PubMed article sets, we generated a dictionary that associates biological entities with key terms that might be used by authors when writing about the biological entity. Biological entities summarize all biological objects and are grouped into 8 biological entity classes: genes and proteins, protein domains, metabolites, diseases, drugs, sub-cellular structures, sub-cellular processes, and confounding terms. Confounding terms were terms that were initially misinterpreted as genes, but referred to an entity that is not part of any of the other entity classes (e.g. “TEP” is an abbreviation for “transepithelial electrical potential”). We generated dictionaries for each of these biological entity classes and combined them to one final dictionary that was used by our text-mining algorithm (Supplementary Fig. S1-12, Supplementary Tables S2-31). The consideration of all biological entity classes helped our algorithm to reduce false positive gene-SCP associations that were based on the misinterpretation of nongene key terms as key terms for genes. We screened the PubMed titles and abstracts of each SCP-specific article set and counted how many articles mentioned a certain gene at least once (Fig. 2). We defined a minimum number of abstracts within an SCP specific abstract that need to mention a gene (level-1 SCPs: at least 4 abstracts, level-2 SCPs at least 3 abstracts, level-3 SCPs at least 2 abstracts, level-4 SCPs at least 1 abstract). Genes that were not mentioned at the specified number of abstracts were removed. Any gene-SCP associations that we labeled as being identified based on the misinterpretation of a non-gene term as gene term during our manual validation process (see below) were removed at this stage. Additionally, we decreased the abstract counts for any gene-SCP association by 66% (an arbitrary selection based on our experience with the population algorithm), if we had labeled this gene during our manual validation to belong to a sibling SCP and not to the SCP it was associated with. These article counts for each gene-SCP association were subjected to statistical enrichment analysis to calculate the selectivity of each gene-SCP association, i.e. how selective is that particular gene for that particular SCP. We used Fisher’s exact test to calculate two different p-values, one with the genes of all SCPs of the same level (same level p-values), and one with the genes of all SCPs that have the same parent and therefore belong to the same children set (same children set p-values) as a background set. Each p-value calculation is preceded by a normalization step: Article counts of each gene-SCP association of a particular process were multiplied with a certain factor, so that after normalization the sum of all article counts of each gene-SCP association of a level-1 and level-2 SCP equaled 3000 and the sum of all article counts of each gene-SCP association of a level-3 and level-4 SCP equaled 1000. This normalization step increased the generation of true positive gene-associations. To calculate p-values we compared the number of articles in a particular SCP specific article set that mention a particular gene with the number of articles in the background set that mention that gene, the total number of articles in the SCP specific article set and the total number of articles in the background set (Supplementary Fig. S16). The resulting p-value estimates the likelihood to observe the documented SCP specific article counts for that gene or a higher one, if it was by chance. We used these p-values to remove un-selective gene-SCP associations. After calculating the negative logarithm of the p-values, we compared the minus log10(p-values) of each association of a particular gene with every SCP in the investigated background set (Supplementary Fig. S17). We determined the largest gap between any two adjacent minus log10(p-values) and defined this gap as the cutoff, i.e. we removed all gene-SCP associations that had a minus log10(p-value) below this gap. Any gene-SCP association that was below this cutoff in the current version of our ontology, but was manually validated as a true positive based on a former version of the ontology was kept nevertheless.

The calculation of the same level p-values depended on a sufficiently high number of SCPs in the background set (>100) that was not given for level-1 as well as on a broad coverage of known cell biology that was not given for level-4 SCPs (not all level 3 SCPs have level-4 children SCPs). Therefore we replaced the filtering based on the same level p-values for these two levels by an inheritance procedure. After finishing the population of level-2 and level-3 SCPs (including manual validation) we only kept those genes in level-1 SCPs that were also part of at least one of its level-2 children or level-3 grand-children SCPs. Similarly, we only kept those genes in the level-4 SCPs that were also part of its level-3 parent. We then proceeded with the calculation of the same children set p-values and filtered the genes as described for level-2 and level-3 SCPs. Finally, we added all genes of level-3 SCPs to their level-2 parents and, following this, all genes of level-2 SCPs to their level-1 parents.

For the manual validation of level-2 and level-3 gene SCP associations (i.e. of those SCPs that were populated based on both enrichment steps), we wrote a script that generates a text file that can be used for a quick validation without any further literature searches. The final text file contains the gene process associations, the description in the NCBI summary of the gene (if available), as well as example sentences describing the gene, protein complex or gene family in the SCP-specific abstract sets (Supplementary Text S1). To allow a visualization of the abstract gene related terms in a different color using a standard text editor (such as JuJu edit in XML mode) we surrounded these terms by greater and smaller signs. Each gene process association was labeled as either a true positive (abbreviation: T), a false positive (F), as identified based on a misinterpreted non-gene term (M) or a false positive that belongs to its sibling process (S). The latter two labels were incorporated into the population pipeline as described above, followed by re-population of the ontology and new manual validation. Since such changes interfered with the population algorithm, and generated new gene-SCP associations, we repeated the manual validation and concurrent re-population of the ontology multiple times until all gene-SCP associations were manually validated. Only true positive associations were kept in the final ontology.

### Analysis of the MBC Ontology

To analyze our ontology, we first documented the number of genes within each SCP and summarized the distribution using the r-functions ‘boxplot’ and ‘beeswarm’. Since level-2 and level-3 SCPs were both populated independently of each other, i.e. based on the same level as well as same children set p-values, we compared the gene composition of level-2 parents and their level-3 children SCPs with each other. This comparison should help us to estimate the consistency of our ontology, since parent SCPs and children SCPs should both be populated with an overlapping set of genes. First, we documented the fraction of genes of each level-3 children SCP that were also part of its level-2 parent SCP (in this case we used a version of the ontology in which parent SCPs have not been populated with the children genes (i.e. step 5 in Fig. 2 was excluded). Results were visualized using R-project. To document, if the annotated parent is the optimal parent, we searched for each level-3 children SCP for that level-2 SCP that contains most of the children SCP genes. The next step was the analysis of how many of the genes of the parent SCP were also part of at least one of its children SCPs (here we included step 5 in Fig. 2). This analysis was repeated after the removal of one or more children SCPs (under consideration of all possible combinations of removed children SCPs) and the complete re-population of all level-3 SCPs (level-2 SCPs were not re-populated). If the children SCPs describe mutually exclusive functions, the removal of any children SCP should decrease the number of genes of the complete children set that overlap with the parent genes. Results were visualized using MATLAB.

For the comparison of our ontology with existing ontologies we downloaded the datasets GO_Biological_Process 2017, Reactome_2016 and Wikipathways_2016 from the EnrichR website^20, 21^.

### Determination of relationships between genes within SCPs by analysis of coexpression data

To test if genes that are part of at least one common SCP show a higher correlation in their gene expression than genes that are not part of any common SCP, we downloaded the median RPKM values by tissue from the GTEx Portal (www.gtexportal.org, file: GTEx_Analysis_v6p_RNA-seq_RNA-SeQCv1.1.8_gene_median_rpkm.gct.gz)^11^. We determined the Pearson correlation coefficient using R-project between any two genes over the 30 tissues. For each level, we considered only those correlations between genes that were both associated with at least one SCP of that level. We separated the correlations into two groups: the first group contained all correlations between any two genes that are part of at least one common SCP of that level, the second group all other considered correlations. Using the function “histogram” in the statistical programming language R we plotted the frequency distribution of the correlations in each group. We used the Two-sample Kolmogorov-Smirnov test (two-sided) in R to determine if the distributions are different.

### Inference of relationships between level-3 SCPs

To predict relationships between level-3 SCPs we re-populated our ontology after removing one level-3 SCP at a time. Here, our population algorithm only removed those manually identified false positive gene-SCP associations that were based on misinterpreted terms; all other false positive associations were kept. The reference ontology that contained the complete set of all SCPs was re-populated based on the same criteria. Nevertheless, we considered only those genes to be members of a left out SCP that were manually validated to be members of that SCP. For each member gene of the left out SCP, we analyzed, if the remaining SCPs contained this gene, and if the final rank of that gene increased because of the removal of the left out SCP. When a gene of the left out SCP became a new member of a remaining SCP, we assumed that its original rank in the remaining SCP was the number of genes of the remaining SCP plus 1. We calculated the sum of all rank decreases within a remaining SCP and normalized it to the size of the SCP before removal of the left out SCP. The normalized sums of each left out remaining SCP combination were used to characterize the strength of the relationship between both SCPs; the higher the sum the stronger the predicted relationship.

### Standard and dynamic enrichment analyses

We used standard enrichment analysis (Fisher's exact test) to analyze experimental published datasets for the enrichment of genes to SCPs in the standard MBC taxonomy and to Gene Ontology Biological Processes (GO_Biological_Process_2017 downloaded from the EnrichR-website). The top 5 predicted level-1, level-2 and level-4 MBC SCPs as well as the top 10 level-3 MBC SCPs were investigated. Calculated p-values were visualized as minus log_10_(p-values) in a bar diagram using R. To visualize parent-child relationships between the top predicted SCPs, we color coded the top predicted SCPs in the annotated MBC hierarchy based on significance. Any ancestors of the predicted SCPs were color coded in gray. All other SCPs were removed from the MBC hierarchy.

For dynamic enrichment analysis, we selected all level-3 SCPs with at least one gene that was perturbed among the experimentally investigated genes and combined up to 3 SCPs if they were related to each other based on the top 25% inferred SCP-SCP relationships between level-3 SCPs (excluding interactions between signaling SCPs, since these most likely arise from shared genes). All possible combinations were considered and added as SCP units to the MBC Ontology (for example, if SCP-A is connected to SPC-B and SCP-B is connected to SCP-C, the SCP combinations A+B, B+C and A+B+C were added as new SCP unions). Experimental gene lists were re-subjected to enrichment analysis using Fisher’s exact test and the extended (context-specific) MBC Ontology. P-values were visualized as minus log_10_(p-values) for the top 5 predicted SCPs or SCP-units. All level-3 SCPs of the top 5 predictions were connected based on all (100%) inferred SCP-SCP relationships (excluding interactions between signaling SCPs). The largest SCP network was identified and manually assigned to a new context-specific higher level SCP. Level-2 parents and level-1 grandparent SCPs were added to demonstrate that the predicted SCPs/SCP-unions belong to different families thus creating an extensible taxonomy.

We tested our ontology on 3 different datasets. Proteomic data^12^ was obtained from supplementary data (Table S3). Genome-wide RNAi screening^13^ results were obtained from supplementary data (Table S4). We used the gene info file that we downloaded from the NCBI website (https://www.ncbi.nlm.nih.gov/gene) to identify the official NCBI gene symbols of the targeted ensembl gene IDs. If an ensembl gene ID matched to more than one official gene symbol, all matches were considered. Gene expression data^14^ was downloaded from GEO repository (GSE30516). Differentially expressed genes were identified using the ‘limma’ r-package.

## Software and tools

C# was used to generate, populate and test the ontology. Matlab and R-project were used to generate Figures. All networks were visualized using yED (https://www.yworks.com/products/yed). Fig. 2, Supplementary Fig. S1-17 as well as the dotted and dashed lines surrounding the SCP-networks in Fig. 6 were made using Microsoft Power Point. For the generation of Supplementary Fig. S15 and S17 we used Microsoft Excel.

## Data Access

Downloaded SCP-specific PubMed abstracts that were used for text mining can be obtained by contacting the corresponding author.

## Acknowledgements

This study was supported by NIH grants GM54508 and P50-GM071558.

## Author contributions

J.H. and R.I. developed the overall concepts for the project. J.H. designed the ontology, developed the algorithms, and tested it. D.M. analyzed co-expression of genes within SCPs. S.W. tested the ontology using standard enrichment analysis. G.S., J.H. and R.I. refined the ontology and J.H., D.M., G.S. and R.I. analyzed the results and wrote the manuscript. R.I. has overall responsibility for this study.

## Additional Information

None

## References

1. Jordan, J. D., Landau, E. M. & Iyengar, R. Signaling networks: the origins of cellular multitasking. Cell 103, 193–200 (2000).

2. Mootha, V. K. et al. PGC-1alpha-responsive genes involved in oxidative phosphorylation are coordinately downregulated in human diabetes. Nature genetics 34, 267–273 (2003).

3. Subramanian, A. et al. Gene set enrichment analysis: a knowledge-based approach for interpreting genome-wide expression profiles. Proceedings of the National Academy of Sciences of the United States of America 102, 15545–15550 (2005).

4. Ashburner, M. et al. Gene ontology: tool for the unification of biology. The Gene Ontology Consortium. Nature genetics 25, 25–29 (2000).

5. Gene Ontology, C. Gene Ontology Consortium: going forward. Nucleic acids research 43, D1049–1056 (2015).

6. Dutkowski, J. et al. A gene ontology inferred from molecular networks. Nature biotechnology 31, 38–45 (2013).

7. Alberts, B., Johnson, A., Lewis, J., Morgan, D. & Raff, M. Molecular Biology of The Cell. Taylor & Francis Group (2015).

8. Berg, J. M., Tymoczko, J. L. & Stryer, L. Biochemistry. New York: W. H. Freeman 5th ed. (2002).

9. Rosenthal, M. D. & Glew, R. H. Medical Biochemistry. John Wiley & Sons, Inc. (2009).

10. Han, J. D. et al. Evidence for dynamically organized modularity in the yeast protein-protein interaction network. Nature 430, 88–93 (2004).

11. Mele, M. et al. Human genomics. The human transcriptome across tissues and individuals. Science 348, 660–665 (2015).

12. Pankow, S. et al. F508 CFTR interactome remodelling promotes rescue of cystic fibrosis. Nature 528, 510–516 (2015).

13. Simpson, J. C. et al. Genome-wide RNAi screening identifies human proteins with a regulatory function in the early secretory pathway. Nature cell biology 14, 764–774 (2012).

14. Lee, M. J. et al. Sequential application of anticancer drugs enhances cell death by rewiring apoptotic signaling networks. Cell 149, 780–794 (2012).

15. Kim, N. & Jinks-Robertson, S. Transcription as a source of genome instability. Nature reviews. Genetics 13, 204–214 (2012).

16. Yang, F., Teves, S. S., Kemp, C. J. & Henikoff, S. Doxorubicin, DNA torsion, and chromatin dynamics. Biochimica et biophysica acta 1845, 84–89 (2014).

17. Engel, N. et al. Actin cytoskeleton reconstitution in MCF-7 breast cancer cells initiated by a native flax root extract. Advancement in Medicinal Plant Research 3, 92–105 (2015).

18. Khatri, P., Sirota, M. & Butte, A. J. Ten years of pathway analysis: current approaches and outstanding challenges. PLoS computational biology 8, e1002375 (2012).

19. Yu, M. K. et al. Translation of Genotype to Phenotype by a Hierarchy of Cell Subsystems. Cell systems 2, 77–88 (2016).

20. Chen, E. Y. et al. Enrichr: interactive and collaborative HTML5 gene list enrichment analysis tool. BMC bioinformatics 14, 128 (2013).

21. Kuleshov, M. V. et al. Enrichr: a comprehensive gene set enrichment analysis web server 2016 update. Nucleic acids research 44, W90–97 (2016).

